# Invariant crossmodal equivalence evokes visual imagery from sounds in rhesus monkeys

**DOI:** 10.1101/2024.01.05.574397

**Authors:** Elizabeth Cabrera-Ruiz, Marlen Alva, Miguel Mata, Mario Treviño, José Vergara, Tonatiuh Figueroa, Javier Perez-Orive, Luis Lemus

## Abstract

After hearing the words Little Red Riding Hood, many humans instantly visualize a girl wearing a red hood in the woods. However, whether nonhuman primates also evoke such visual imagery from sounds remains an open question. We explored this from direct behavioral measurements from two rhesus macaques trained in a delayed crossmodal equivalence task. In each trial, they listened to a sound, such as a monkey vocalization or a word, and three seconds later, selected a visual equivalent out of a pool of 2 to 4 pictures appearing on a touchscreen. We show that monkeys can be trained to discriminate perceptual objects of numerous properties and furthermore that they perceive as invariant different versions of the learned sounds. We propose two potential mechanisms for the brain to solve this task: acoustic memory or visual imagery. After analyzing the monkeys’ choice accuracies and reaction times in the task, we find that they experience visual imagery when listening to sounds. Therefore, the ability of rhesus monkeys to perceive crossmodal equivalences between learned categories poses rhesus monkeys as an ideal model organism for studying high-order cognitive processes like semantics and conceptual thinking at the single-neuron level.

## Introduction

Crossmodal correspondence, a natural animal ability, ‘translates’ information from one sensory modality to another (Parise et al., 2016), for example by taking advantage of how they both vary in a single physical dimension, e.g., lightness, roundness, texture, loudness, and pitch (Walker et al., 2012; Ratcliffe et al., 2016). In lab settings, chimpanzees discriminate visual shapes using sounds (Ravignani and Sonnweber, 2017), and trained macaques perform crossmodal discriminations of shapes and textures (Weiskrantz and Cowey, 1975; Zhou and Fuster, 2000), tones and colors (Fuster et al., 2000), and flutter frequencies (Lemus et al., 2010).

However, all these studies explored simple crossmodal correspondence along one physical dimension such as shape or color, whereas in real life, we know that humans form complex crossmodal equivalence (CME) between items of multiple and heterogeneous physical attributes, for example, between words and the visual concept they evoke. Visualizing ideas from words is a complex ability, facilitating the execution of intricate tasks, such as building a cathedral. However, such a crucial evolutionary trait may have appeared first in an ancestor common to other nonhuman primates (NHP). For instance, ethological studies implicitly suggest CME in NHP (Ratcliffe et al., 2016). In a classical paper by Seyfarth and colleagues (Seyfarth et al., 1980), vervet monkeys possibly evoked visual imagery because they looked up, down, or ran into the trees after hearing conspecific alerts of leopards, eagles, and snakes.

Neuroanatomical and behavioral similarities between humans and NHP indirectly suggest that NHP can experience CME, for example, between voices and faces (Jordan et al., 2005; Adachi and Hampton, 2011; Tyree et al., 2023). This mechanism could rely on the integration of visual and auditory representations in multisensory areas of the brain (Gaffan and Harrison, 1991; Calvert et al., 2001; Beauchamp et al., 2004; Khandhadia et al., 2021; Diehl et al., 2022; Lemus and Lafuente, 2022), or from connecting neuronal representations of monkey calls at the superior temporal gyrus (STG) (Leaver and Rauschecker, 2010; Tsunada et al., 2011; Bodin and Belin, 2020; Bodin et al., 2021) with unitary activity associated to a monkey face at the superior temporal sulcus (STS) (Tsao et al., 2003; Leopold et al., 2006; Ohayon et al., 2012; Arcaro et al., 2017; Khandhadia et al., 2021). Regardless of NHP being able to discriminate words phonetically (Melchor et al., 2021), it is unclear whether their brains can encode complex sounds like words (Hickok and Poeppel, 2007; Yi et al., 2019; Morán et al., 2021; Stephen et al., 2023) and associate them with visual representations to experience CME.

Therefore, in the present study, we explore the ability of monkeys to perceive CME between sounds, including words, and pictures rather than just triggering stereotyped behaviors. Thus, we trained two rhesus macaques in a delayed CME task. In each trial, a reference sound (S) appeared, and three seconds later, the monkeys had to match the sound with its equivalent picture (P) from a pool of pictures (Pics) presented on a touchscreen. Different hypotheses have been proposed about the perception of an object from different modalities (Walker et al., 2012; Parise et al., 2016). We test two potential models for solving the task: In the first, S elicits visual imagery in working memory to compare against Pics. In the second, the S identity persists in auditory working memory. Our results strongly suggest that the monkeys experienced visual imagery during the CME task.

## Results

We trained two rhesus monkeys in a CME task. In each trial, the monkeys listened to an S and waited 3 seconds to find the corresponding P out of several Pics (**Fig 1A**, **S1 Video 1**; see methods). Solving the task required the monkeys to hold information regarding S in working memory. We hypothesized that such information would consist of visual imagery of P rather than a trace of S. In other words, monkeys had to ‘translate’ S into P to discriminate Pics in the same sensory modality. Therefore, we sought to find whether evoking learned visual information about P predicted the animals’ choice accuracy and reaction times (RT).

**Fig 1.**
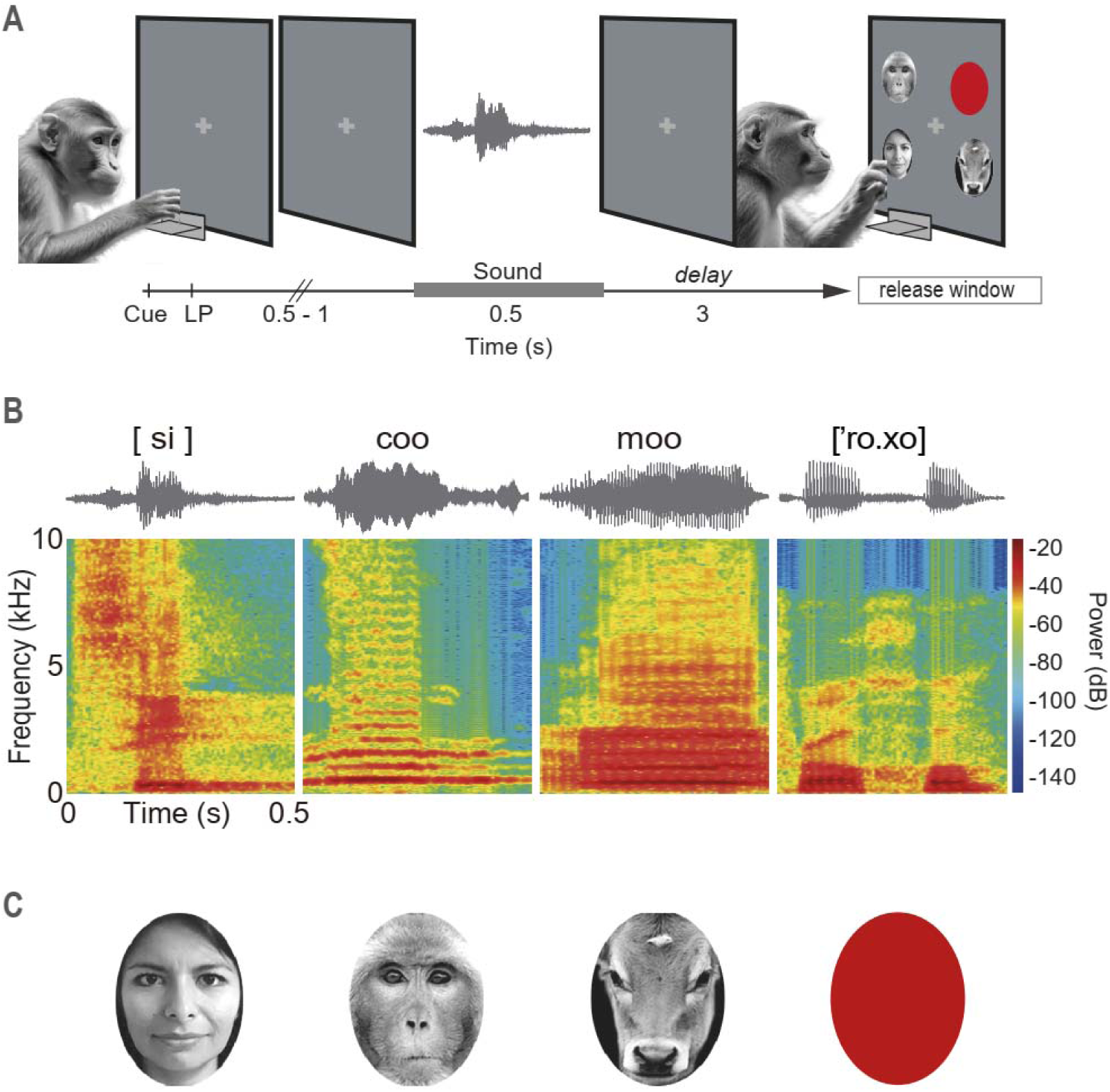
Delayed crossmodal equivalence task. **(A)** Task events. In each trial, the monkey pressed a lever in response to a grey cross on the center of the touchscreen. After a variable period of 0.5-1 s, a reference 0.5 s S was delivered. Then, after a 3 s delay, several Pics were presented simultaneously on the touchscreen, and the animal released the lever to select the P equivalent to S. Correct responses were rewarded with a drop of liquid. LP, lever press. **(B)** Sonograms and spectrograms of some S in the task, including two Spanish words shown in IPA nomenclature. Other sounds corresponded to a monkey coo call and cow vocalization. **(C)** Pictures learned as equivalents of the S shown above in **B**. Each CC consisted of an S-P pair.

## The monkeys learned crossmodal equivalences in a few sessions

We created eighteen crossmodal categories (CC) from combinations of six S and fourteen P (**S2 Table of CCs**; See examples in **Fig 1B and C**). The monkeys initially practiced the task in trials with only two Pics, one equivalent and one nonequivalent (NE; see methods) to S. Then, we introduced more CCs and trials of 3 and 4 Pics. To analyze the monkeys’ learning process, we derived four learning parameters from fitting simple associative learning curves to the performance in each CC throughout sessions: the hit rate (HR) in the first session (y0) and sessions of statistical (y), inertial (*δ*), and consolidated learning (λ), respectively (see Methods).

The performance of monkey G at three CCs discriminated in trials of 2, 3, and 4 Pics is shown in **Fig 2A**. Since the coo/monkey-face CC (left panel) was one of the first learned, its y0, or initial performance (∼300 trials) was close to chance level (intersection of the black learning curve with the ordinate). Then, it took another eight sessions (∼2700 trials) to reach statistical learning (y; **Fig 2A**, gray boxes, left edge) when the subjects began to solve the task without being consistently correct. Then, inertial learning (*δ*) occurred after the 15^th^ session, where the change in performance from one session to another was insignificant, so the performance is assumed to be constant. Our interpretation is that a transition from sufficient performance (y) to efficient (*δ*) behavior occurred before reaching the optimal performance (λ).

**Fig 2.**
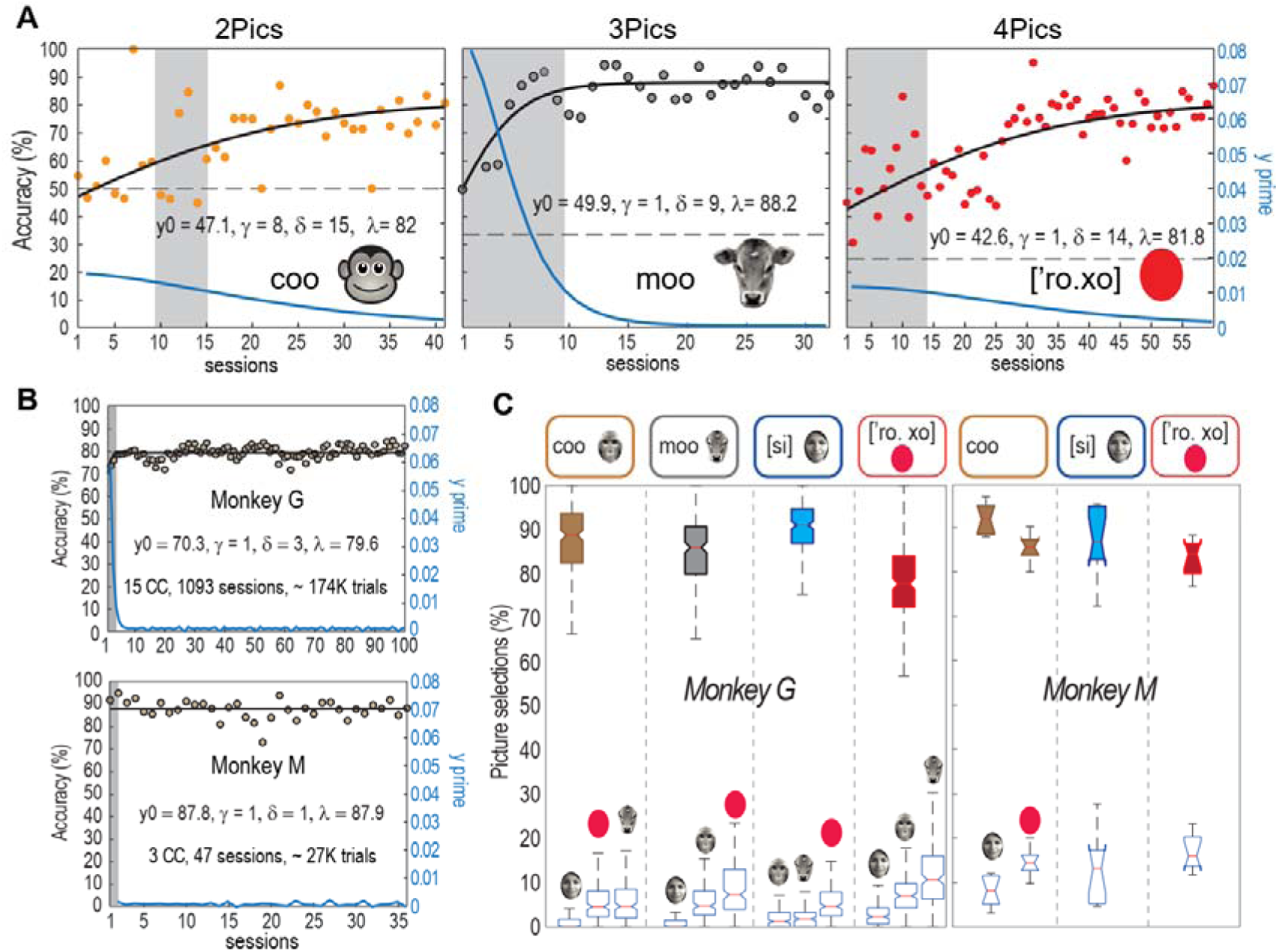
Monkeys learn the crossmodal equivalence between sounds and pictures. **(A)** Sigmoidal fits (black lines) of the behavioral accuracy of monkey G throughout learning sessions. Blue lines, y’ values of the first derivative of the sigmoidal fits. Each panel corresponds to the accuracies in the CCs monkey, cow, and [‘ro. xo] (the Spanish word for red), discriminated in trials of 2, 3, and 4 Pics on the touchscreen, respectively. Grey boxes delimit the sessions between the statistical learning y and the inertial learning o (see methods). Dashed lines indicate the chance level at each number of Pics. (**B)** Same as in **A**, but for the averaged accuracy of all CC performed by each monkey. (**C)** Boxplots of the overall proportion of hits (closed) and false alarms (open) of sessions after o.

The number of sessions for reaching y and o at new CCs decreased when the monkeys learned the aim of the task. For example, monkey G took less than three sessions to reach y in trials of 3 and 4 Pics and less than ten to o. The exception was the word [ro. xo] (Spanish for red), where it spent ∼14 sessions to o. **S2 Table of CCs** shows all CCs and their learning parameters. **Fig 2B** presents the mean of all CC learning for both monkeys. We interpret the reduction in y and o as the monkeys solving the cognitive control of the motor behavior required for the task (procedure memory), e.g., pressing and releasing the lever and interacting correctly with the touchscreen, so that once this was done, the animals could focus only on learning the CC associations.

**Fig 2C** shows examples of boxplots of the monkeys’ overall accuracy after o sessions, i.e., after approaching the asymptote of learning the CC. In the case of monkey G, CCs were presented in trials ranging from 2 to 4 Pics, whereas for monkey M, with 2 Pics (**S3 Table Overall performance**). The open boxplots at the bottom of the figure correspond to false alarms (FA), estimated as the percentage of times nonequivalent Pics were selected. The overall HR of monkey G was 85.12 ± 9.11 (mean ± SD), and for monkey M, 87.07± 5.71 (mean ± SD). We also evaluated the performance of each category as a function of the angles at which the correct visual cue was presented and found no bias (**S4 Figure)**. Regardless of the low proportion of FAs, they differed with every P so that, for each CC, there was a sequence in FA distributions. For example, for monkey G, there often was a higher proportion of FA for the picture ‘red,’ then for the cow, followed by the monkey, and, finally, the human, suggesting a relationship between FAs and visual information, which we explore further in the following sections. These results demonstrate that the monkeys achieved the CME task proficiently after a few training sessions and with discriminations of 2 to 4 Pics.

## The RT distributions correlate with the number of visual categories

We then explored the impact of Pics and each P on the reaction times (RT, from the appearance of Pics to lever release) and motor times (MT, from lever release to the touch of a P). **Fig 3A** shows that while the MT are similar for all 4 CCs, the RT clearly differentiate between CCs, with the ‘human’ (H) CC having the longest RT. Furthermore, S having two syllables as in [‘ro. xo] yielded the shortest RT, suggesting that the number of syllables of the auditory stimuli is not directly linked to RT. Similarly, for all CCs, the higher the mean RT, the higher the variance (**Fig 3C**, top). Notably, no correlation existed between HR and RTs (**Fig 3C**, bottom, Spearman rank correlation test).

**Fig 3.**
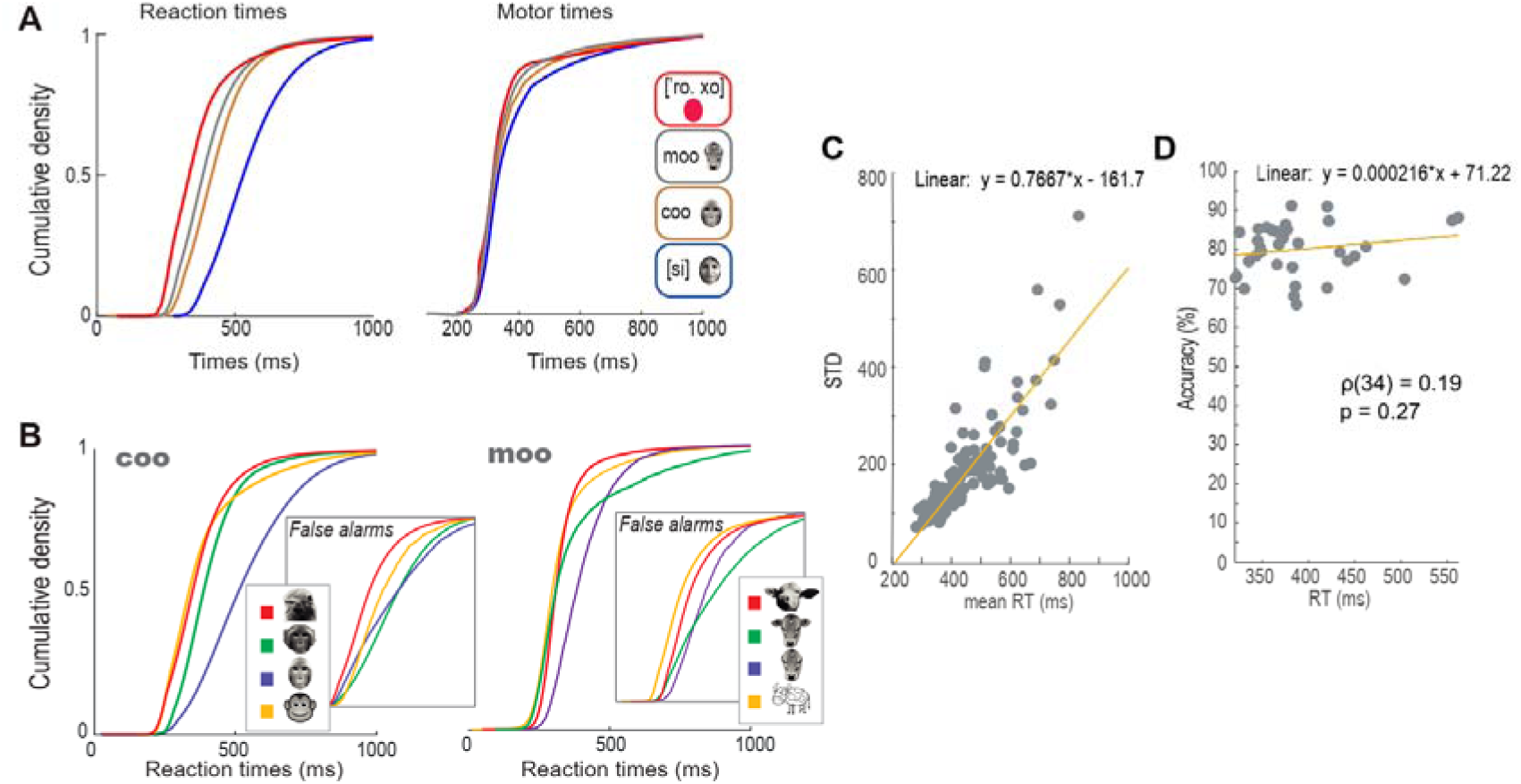
RTs increase as a function of Pics. **(A)** Cumulative density functions for RT (left) and MT (right) for monkey G. **(B)** Cumulative density functions of RTs in various P, all equivalent to a single S. Each panel shows RT distributions of correct responses to different P, all equivalent to a constant S. Insets show RT of FAs produced when each P was selected, while presenting a nonequivalent S. Notice how RT distributions resemble hits so that P selections rather than S seem to bias RT. (**C)** Standard deviation of RT distributions as a function of the distributions’ mean (top); and relationship between RT and accuracy (bottom); RT did not affect accuracy.

By analyzing the RTs of different P in monkey G, which are all equivalent to a single sound (e.g., four monkey faces equivalent to the same coo vocalization) (**Fig 3B**), we found that the shape of the RT distributions were different for each P (p < 0.001, Kruskal-Wallis test, p < 0.001 for all pairs of P and same S, post hoc Mann-Whitney U test, Bonferroni corrected).

Therefore, given that the S was the same, the differences in the RTs are due to the P categories. The same happened in FA (**Fig 3B, insets**), where each distribution corresponded to the selection of a P nonequivalent to several sounds (p < 0.001, Kruskal-Wallis test, p < 0.01 for 71.43% for pairwise comparisons of coo, 76.19% for moo, and 82.14% for [si], Mann-Whitney U test, Bonferroni correction). In other words, the RT distributions changed with P visual stimuli while keeping S constant.

## The monkeys showed auditory perceptual invariance and RTs that followed the visual categories

So far, we have inferred that changes in RTs are not explained by S because they change while keeping S constant. To further test this, we presented the monkeys with versions of S from emitters to which they had no previous exposure (**Fig 4A**). **Fig 4B** shows performance above chance in monkey G in 98.33% of the cases (paired-sample t-test, p < 0.05; see **S6 Table Versions**). Notably, the RTs of hits at most versions were dispersed in P visual categories rather than S auditory versions (**Fig 4C**).

**Fig 4.**
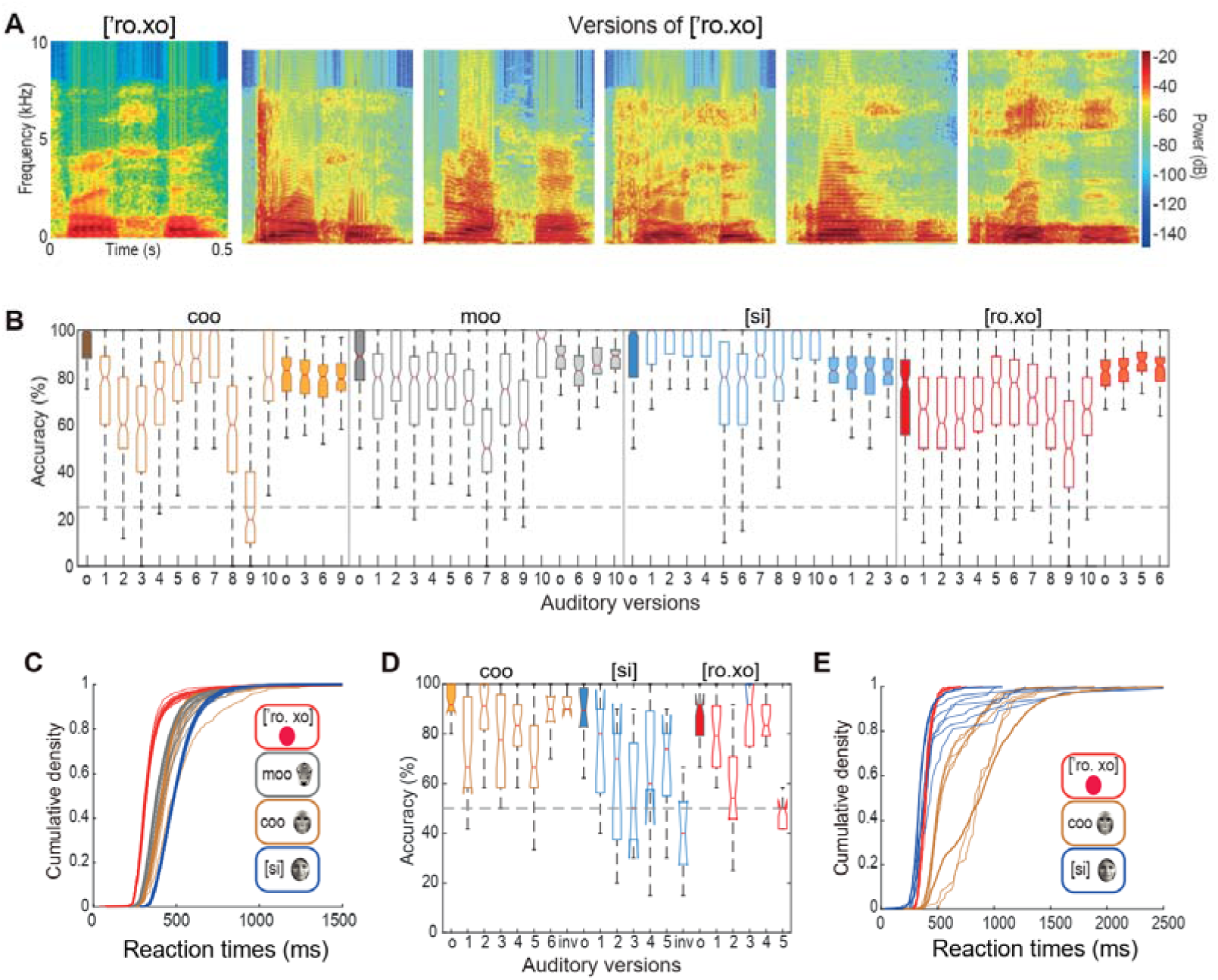
Monkeys show perceptual invariance to versions of S, reacting in times of P categories. **(A)** Spectrograms of some versions from different emitters of [‘ro. xo] word. **(B)** Boxplots of Monkey G accuracies with all S versions. Each color corresponds to each S category. Closed boxplots at the left of each category correspond to the accuracy during the learned sound (o). Open boxplots correspond to versions of o. Closed boxplots at the right of each group correspond to versions which were repeated twice within 0.5 seconds. **(C)** RTs of hits of versions in **B,** remain close to o’s RTs (bold lines) and are distributed into the number of P categories. **(D)** Same as **B**, but for monkey M. **(E)** Same as **C**, but for monkey M.

Meanwhile, monkey M performing in trials of 2 Pics, showed performance above chance at 72.22 % of versions (5 to 7 versions per sound; paired-sample t-test., **Fig 4D**). Here, RTs for hits and FAs were also dispersed in P categories (**Fig 4E**). These results suggest that the monkeys experienced auditory constancy, leaving visual imagery as the prevailing explanation for varying RTs during the CME task. Thus, we found that monkeys could generalize, assigning auditory stimuli from different emitters to the learned S category. Also, the RTs from the various emitters are explained by visual but not auditory properties.

## DISCUSSION

In our study, we trained two rhesus monkeys to perform a delayed crossmodal equivalence task to recognize picture equivalents of sounds. We report three main findings from the testing of the animals in different conditions. First, the monkeys can learn in a laboratory setting crossmodal associations between visual and auditory objects as complex as faces, cartoons, and words (**Fig 2**). Second, they perceive as invariant different versions of the learned sounds (**Fig 4**), and third, sounds evoke visual representations in their minds (**Figs 3-4**).

### Rhesus macaques learn crossmodal equivalences across complex heterogenous stimuli

In contrast to studies demonstrating the ability of monkeys to discriminate bimodally auditory, visual, or tactile stimuli with clear correspondence along one physical dimension (Weiskrantz and Cowey, 1975; Zhou and Fuster, 2000; Lemus et al., 2010; Nieder, 2012), for instance, the frequency of a tone with the frequency of a tactile vibration (Lemus et al., 2010), we show that they can be trained in a laboratory setting to discriminate perceptual objects of numerous and heterogeneous properties, as humans do between words and images. This is consistent with evidence that neurons in the monkey’s hippocampus respond invariantly to the face and voice of conspecifics in the colony (Tyree et al., 2023).

Monkeys have consistently proven their capabilities in bimodal discrimination tasks (Gaffan and Harrison, 1991; Fuster et al., 2000). However, the mere training of monkeys in auditory tasks has been considered challenging (Ng et al., 2009; Scott et al., 2012). We also found it difficult since when we initially paired visual and auditory stimuli, the monkeys ignored the sounds. Thus, we trained them to detect the sounds first and do crossmodal associations afterwards. Once they grasped the task’s objective, the learning time of new CC categories was reduced significantly. Finally, we derived novel behavioral parameters from the performance throughout sessions to understand their learning process. The parameters, such as statistical and inertial learning derived from sigmoidal fits to performance, could help tailor individual monkey training.

### Learned sounds evoke visual imagery in monkeys

It has previously been reported that animals can imagine (Lai et al., 2023). However, whether sounds evoke visual imagery in NHPs or set off stereotyped behaviors is unclear. We present three results suggesting that sounds evoke visual imagery in monkeys. First, reaction times in the CME task differentiate between visual, not auditory stimuli (**Fig 3**). Second, because of the monkeys’ auditory perceptual invariance, the RTs did not disperse into new acoustic categories but remained in distributions of visual categories (**Fig 4**). Consequently, stereotyped reactions do not explain the monkeys’ behavior. Instead, their decisions rely on the discrimination of visual images evoked by the sounds.

Additionally, the bi-syllabic words [’ro. xo] and [’t[aŋ.gi], carry more auditory information than the coo and the moo vocalizations. Therefore, in the auditory working memory scenario, the slowest RTs should have correlated with those words, but this was not the case. We further assessed the effect of sound repetitions, e.g., coo-coo and [si]-[si], finding that the performance was faster and more accurate than in single presentations, and again, the RTs were grouped into visual categories. Another clue was that both monkeys reacted faster to the “red” category, which consisted of only one visual attribute (even though it is a bi-syllabic word in Spanish). At the same time, they spent more time discriminating the monkey and human faces with similar attributes. In conclusion, the more complex the visual discriminations, the longer it takes to achieve CME.

Similarly, sounds from different emitters are dispersed in a few visual categories rather than in the numerous acoustic versions. In other words, like humans, monkeys experience auditory perceptual invariance (Town et al., 2018) and make general relationships with visual information. Although we only presented a limited number of visual equivalents, visual imagery likely emerges during the delay period because, as suggested by previous studies, discriminating auditory information is hard for macaques (Ng et al., 2009; Scott et al., 2012), so visual imagery would present more naturally.

Multisensory areas like the prefrontal cortex or the STS, which store visual objects (Beauchamp et al., 2004) and receive auditory inputs (Noesselt et al., 2007), are likely to integrate acoustic and visual objects. We envisioned two alternative mechanisms to achieve the CME: acoustic memory and visual imagery models. In the former, a sound would activate its putative neural circuit at STG and remain active in working memory (**Fig 5A**). Then, when its visual equivalent activates a visual representation at STS, it increases the STG firing rate via a direct input to reach a CME threshold (**Fig 5B**). This model implies that crossmodal equivalence translation only occurs during the comparison period.

**Fig 5.**
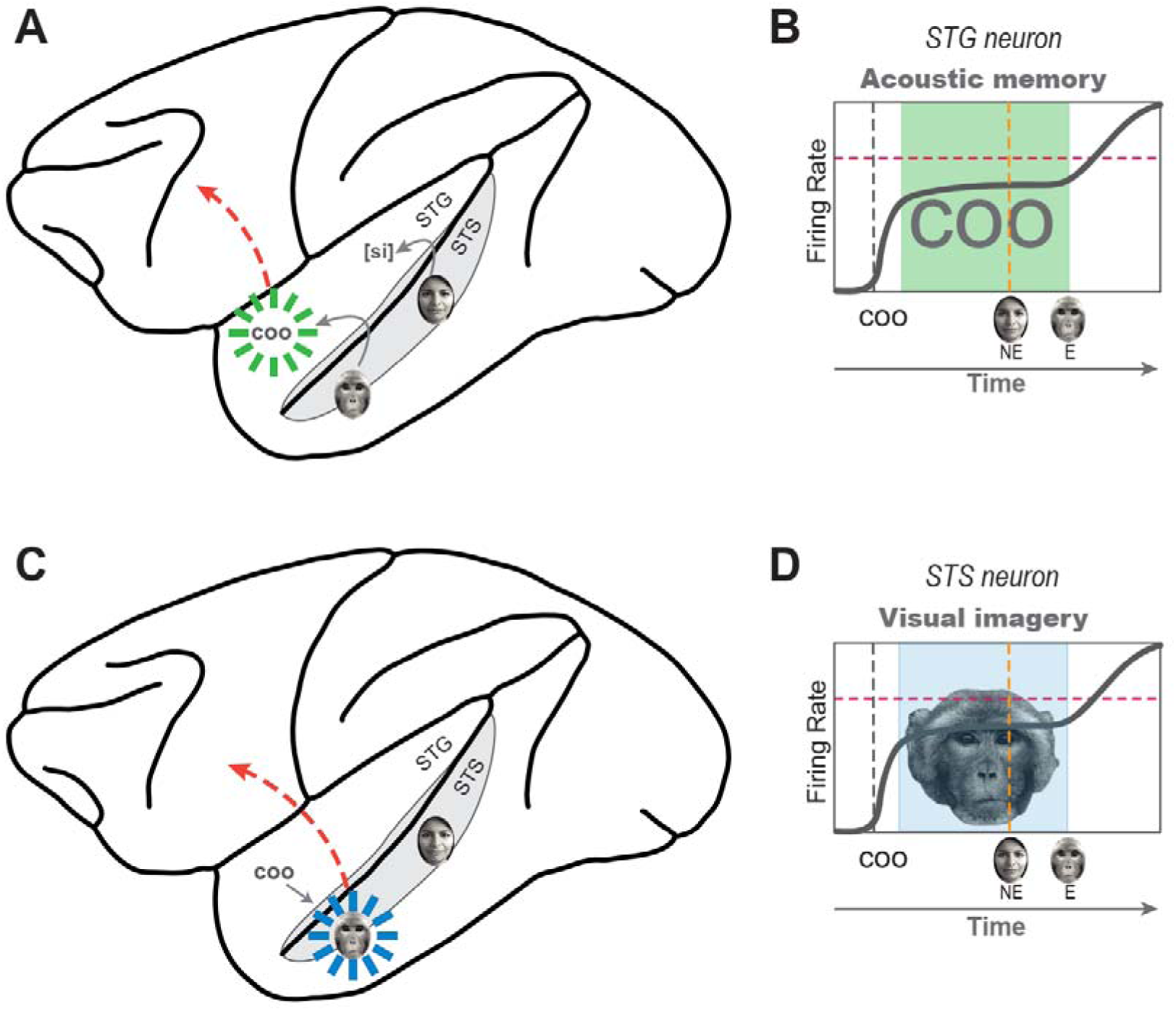
Models for delayed CME in the brain. **(A)** Auditory working memory model. When a sound elicits an STG circuit, it remains active until a signal of the visual equivalent evoked at STS arrives to trigger a perceptual equivalence to send to executive brain areas (dashed red arrows). **(B)** Firing rate of an STG neuron across events of the CME task (bold line). The neuron responds to a sound (gray dashed line) and remains active during the delay (green box). Since the neuron has no inputs of nonequivalent visuals (NE; yellow dashed line), its firing rate remains constant. However, when the equivalent P activates STG through direct input from STS, the firing rate increases above the perceptual equivalence threshold (red dashed line). **(C)** Visual imagery model. The arrival of a sound activates STG, which activates STS, evoking visual imagery of P. Visual imagery remains until the actual sight of P creates perceptual equivalence, which is then communicated to executive brain areas (dashed red arrow). Notice that NE P has no interaction with the visual circuit activated by STG. **(D)** Same as in **B**, but for a neuron in STS and visual imagery active in working memory.

Alternatively, in the visual imagery model (**Fig 5C**), once an STG circuit gets activated by a sound, it immediately activates its visual equivalent at STS, which remains active as visual imagery during the delay period until a picture on the touchscreen increases its firing rate to equivalence threshold (**Fig 5D**). This model would explain the visual imagery evoked from sounds. Our experiments demonstrate that the reaction times correlate with evaluating visual information. Future recordings of the neural responses in areas such as STG and STS can provide essential additional information (Cohen et al., 2007; Romanski, 2007; Chandrasekaran et al., 2013; Huang and Brosch, 2016; Diehl et al., 2022).

Our findings prove that monkeys visualize images upon listening to communication sounds. This crossmodal ability may have evolved in humans and monkeys from a common ancestor to identify predators or rivals and understand other monkeys’ emotional states (Bodin et al., 2021; Diehl et al., 2022). Based on our results, we propose that macaques present an ideal model for studying the neural basis of crossmodal equivalence. This phenomenon is probably involved in other cognitive processes, such as the semantic process in language and conceptual thinking.

## Methods

### Ethics statement

All experiments observed the Official Mexican Standard Recommendations for the Care and Use of Laboratory Animals (NOM-062-ZOO-1999) and were approved by the Internal Committee for the Use and Care of Laboratory Animals, Cell Physiology Institute, UNAM (CICUAL, LLS200-2).

### Subjects

Two adult rhesus monkeys (*Macaca mulatta*) participated in the experiments (a female, monkey G, 10yr, 7 Kg, and a male, monkey M, 12yr, 12 kg). The animals inhabited single cages in a room with constant temperature (22°C), filtered air, and day/light transitioning lights. The animals had access to sources of enrichment, such as TV, toys, a recreating area, and grooming with other animals through mesh sliding doors. They were restricted to water for 12-15 hr. before experiments (Monday-Friday water intake > 20-30 ml/kg, ad libitum on weekends). In addition to the primate pellets, they received 150 mg rations of fruits and vegetables after the 2-3 hr. experimental sessions.

### Experimental setup

The primates were trained to leave their cages and sit on a primate chair (Crist Instrument, INC.) to be transferred to a soundproof booth adjacent to the vivarium for experiments.

Then, the chair was placed 30 cm from a touchscreen (ELO 2201L LED Display E107766, HD wide-aspect ratio 22in LCD) and a spring lever below the touchscreen (ENV-610M, Med Associates). Two speakers (Yamaha MSP5 Studio, 40 W, 0.050-40 kHz, and Logitech 12 W, 0.01-20 kHz) above the touchscreen delivered the sounds and a background noise at 45– and 65-dB SPL, respectively. The monkeys obtained liquid rewards through a stainless-steel mouthpiece attached to the chair (Reward delivery system 5-RLD-E2-C Gravity feed dispenser, Crist Instrument INC.).

### Acoustic Stimuli

Sounds consisted of lab recordings of words, monkey vocalizations, and sounds downloaded from free online libraries. All sounds were resized to 500ms, resampled to 44.1 kHz (cutoff frequencies 0.1–20 kHz), and finally, normalized (RMS; Adobe Audition® 6.0). For the perceptual invariance experiment, we replaced the sounds with versions of the same categories and durations but with different emitters. Phonetic nomenclature of words in Spanish was obtained using the Automatic Phonetic Transcriptionist by Xavier López Morrás (http://aucel.com/pln/transbase.html).

### Visual Stimuli

The visual stimuli consisted of black and white photographs of cows, monkeys, and human faces, two cartoons, and a red oval circumscribed in ovals (200px/ sq-inch resolution), presented equidistant from each other within a 4° radius from the center of the touchscreen. The human face corresponds to the portray of the first author in this manuscript. Other pictures were downloaded from free online sites and customize for the porpoise of the experiments. Artwork of monkey in **Fig 1A** was created using a free online platform (https://www.fotor.com/ai-art-generator).

### Delayed crossmodal equivalence task

We trained two rhesus macaques in a delayed CME task. Each trial commenced with a 2° aperture gray cross on the center of the touchscreen, indicating the monkey to press and hold down the lever. Then, a 0.5 s sound followed by a 3 s delay period was delivered. Finally, a pool of 2-4 simultaneous pictures appeared on the touchscreen, so the monkeys had to release the lever and select the visual equivalent within a 3 s response window. The animals obtained a drop of liquid reward for correct selections. Sets of trials comprised ∼300 trials of 2-4 CC presented intermingled. The task was programmed in LabVIEW 2014 (64-bit SP1, National Instruments®).

### Monkeys training

Our primary objective was to get the monkeys to perform the CME task as quickly and efficiently as possible. Therefore, instructions included changes in stimuli, durations, and rewards, sometimes within a single session, depending on the ongoing behavior of the monkeys. Our approach was to train the animals to respond to a visual cue by pressing the lever and interacting correctly with the touchscreen. Afterward, we introduced trials starting with a sound and ending with two pictures. Only after they learned the first two CCs, we presented the rest.

### Behavioral measurements

Although our experiments did not aim to study the learning process of macaques, we observed behavioral improvements across the sessions we sought to describe. First, we fit learning curves to performance at each CC across sessions to evaluate the monkeys’ learning process. To establish a criterion for identifying discriminative choices within steady-state behavior, we fitted learning curves to the monkeys’ choice records using the Rescorla– Wagner model of associative learning (Rescorla and Wagner, 1972). This model conceptualizes learning through associations between conditioned and unconditioned stimuli. The extraction of the learning curves involved numerically solving the following ordinary differential equation, as described before (Treviño, 2016):

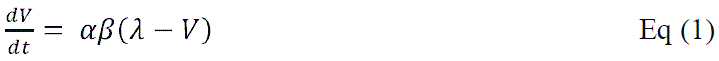

This equation describes the evolution of the associative strength (*V*) towards a trained conditioned stimuli as a function of the training trials (*t*). Optimized parameters from the model were αβ, the product of the salience of the conditioned stimuli and the strength of the unconditioned stimuli (which were assumed to be fixed during training, but see (Treviño, 2016), and the asymptote of learning (A), which corresponds to the maximum conditioning possible for the unconditioned stimuli. We then derived three additional parameters from the extracted learning curves. Y0 described performance during the first training session, corresponding to the fit intersection with the ordinates. Then, y, or statistical learning, was the first session at which performance statistically differed from chance, i.e., it surpassed two standard deviations from chance level in a binomial distribution, where p = chance level and n = mean number of trials per session. Meanwhile, the first derivatives of the extracted learning curves were combined with user-defined thresholds to identify the inertial (*δ*) experimental session at each CC where the monkey choices approached the A learning plateau, i.e., when the change in performance between two consecutive sessions was equal or less than arbitrarily selected threshold of y’ = 0.01 (Figure 2A, blue lines).

Reaction times (RTs) were the times from the appearance of the pictures to the lever release. Choice accuracy was defined as the proportion of correct responses. False alarms consisted of selections of nonequivalent pictures. Early responses and misses (not included in this study) consisted of lever releases before or after the response window, leading to aborting the trials.

To analyze biases toward a position on the screen for a specific P category, we analyzed the difference between angles using the following quadrant approach. For a given P angle (reference angle), we defined its reference 90°-quadrant as the area 45 degrees to the left and 45 degrees to the right of that angle. We drew the rest of the 90°-quadrants relative to the reference one. Then, using the multiple pairwise comparisons’ results from the one-way ANOVA (Tukey’s HSD, p<0.05), we classified a quadrant as significant if at least half of its angles significantly differed from the reference angle.

### Statistical analyses

Most analyses in this study comprised sessions after the *δ* session. Monkey M performed in trials of two Pics and monkey G four Pics. Thus, performance above chance was greater than 0.5 for monkey M and 0.25 for monkey G. We only included RTs from sessions after o performance to compare RT distributions. We performed Spearman rank correlation tests to find statistical differences between CCs RT distributions as a function of the number of Pics. We performed a Kruskal-Wallis test to assess statistical differences among crossmodal categories. In the case of null hypothesis rejection, a Mann-Whitney test was used to compare cases of 2 and 3 P equivalent to one S and cases of a learned S with its versions. A Bonferroni post hoc test was performed for multiple comparisons. Analyses were performed using MATLAB R2022 (MathWorks).

### Author contributions

Conceptualization: LL. Data Curation: ECR, MM, TF. Formal Analysis: ECR, MA, MT, MM, JV & LL. Funding Acquisition: LL. Investigation: ECR, MM. Methodology: ECR & LL. Software: ECR, MA, TF, MT, & LL. Validation: JPO, MT, JV & LL. Visualization: ECR, MA & LL. Writing original draft: LL. Review & draft editing: ECR, MA, JPO, MT & JV.

## Abbreviations

CC: Crossmodal categories
CME: Crossmodal equivalence
FA: False alarms
HR: Hit rate
NE: Nonequivalent
NHP: Nonhuman primates
MT: Motor times
P: Picture
Pics: Pool of pictures on the touchscreen
RT: Reaction times
S: Sound
STG: Superior temporal gyrus
STS: Superior temporal sulcus

## Acknowledgments

We thank Francisco Pérez, Gerardo Coello, and Ana María Escalante of the computing department of the IFC, Aurey Galván and Manuel Ortínez of the IFC workshop, and Centenario 107 for providing invaluable working space and edible resources. Elizabeth Cabrera Ruiz conducted this study to fulfill the requirements of Programa de Doctorado en Ciencias Biomédicas of Universidad Nacional Autónoma de México and received a doctoral scholarship from Consejo Nacional de Humanidades Ciencias y Tecnologías (245771). The data in this work are part of her doctoral dissertation.

## Supporting Information

**S1 Video 1.** Monkey G performing in the delayed CME task.

**Table 1.**
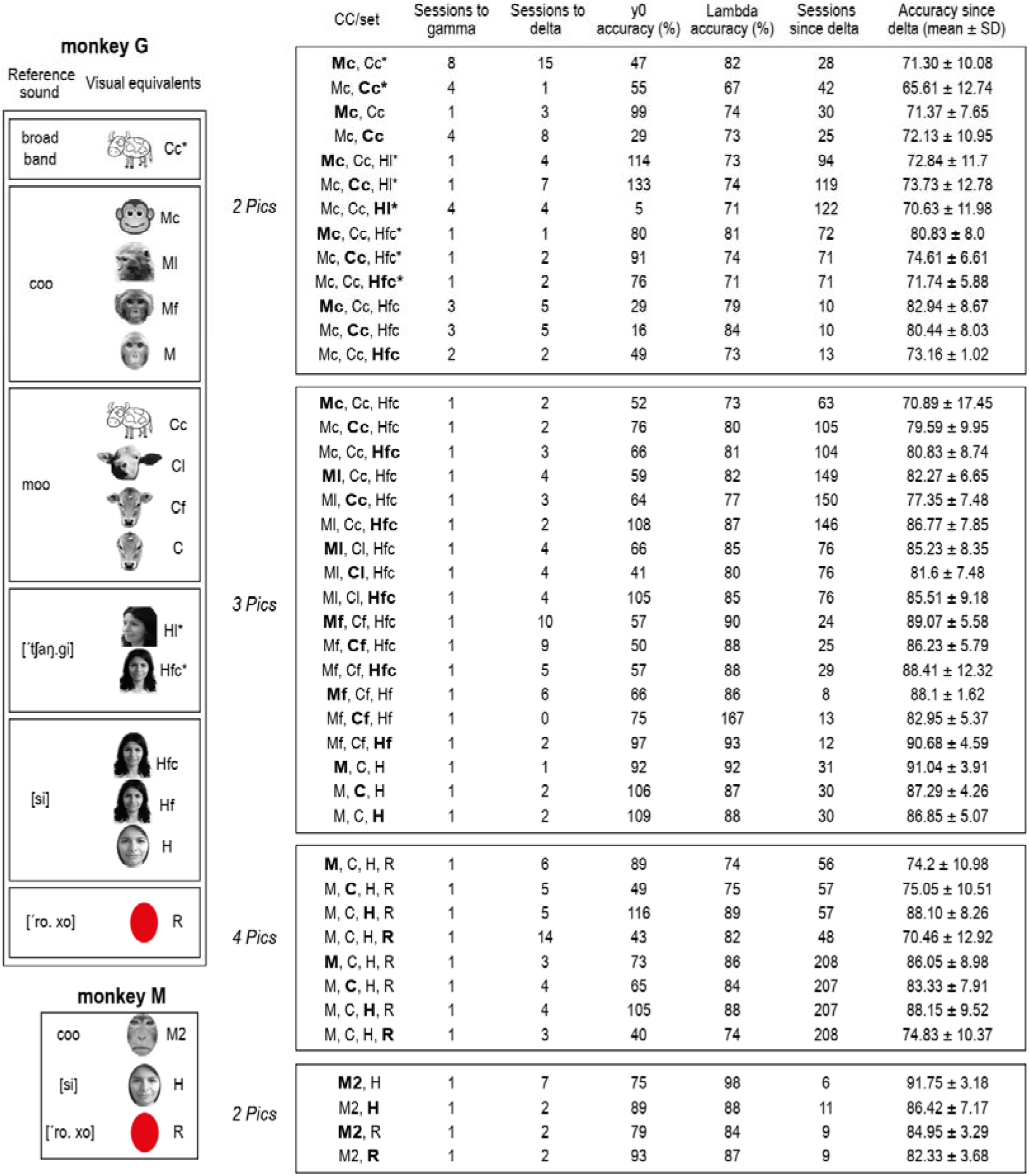
Monkey’s accuracy in crossmodal categories. **S2 Table of CCs.** Table presenting all crossmodal categories and the monkeys’ parameters obtained from their learning processes

**Table 2.**
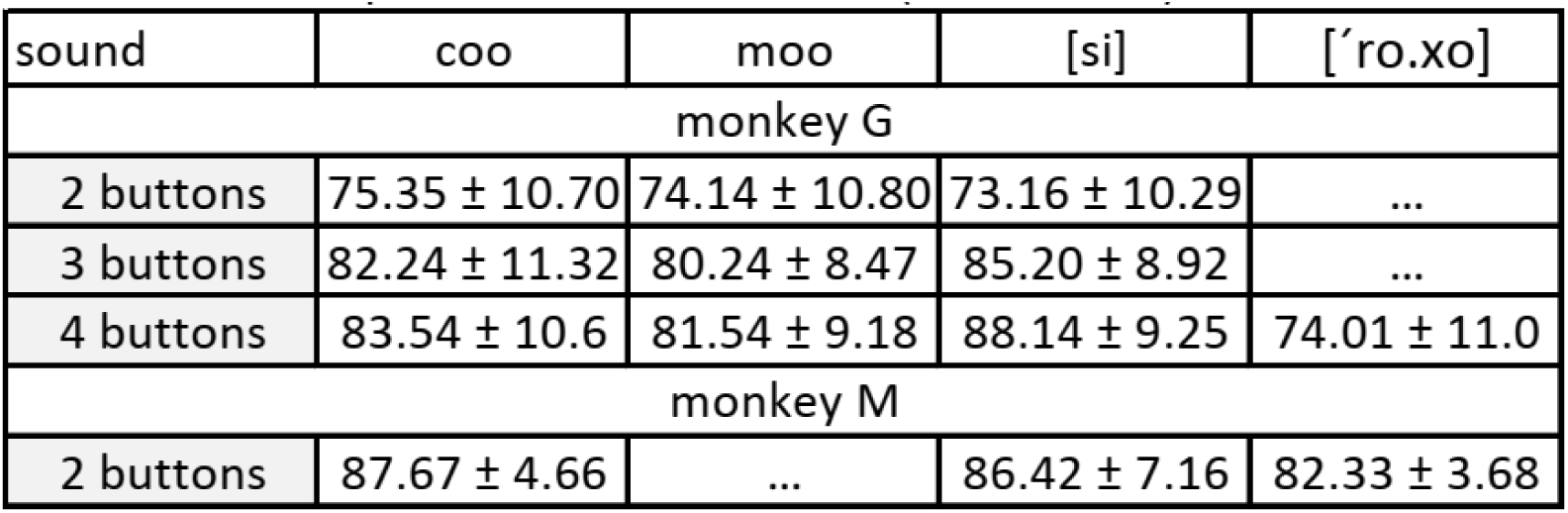
Overall performance after delta (mean ± s.d.) **S3 Table Overall performance**. Monkeys’ performance in CCs most used.

| sound | coo | moo | [si] | [‘ro.xo] |
| --- | --- | --- | --- | --- |
| monkey G |  |  |  |  |
| 2 buttons | 75.35 ± 10.70 | 74.14 ± 10.80 | 73.16 ± 10.29 | ... |
| 3 buttons | 82.24 ± 11.32 | 80.24 ± 8.47 | 85.20 ± 8.92 | ... |
| 4 buttons | 83.54 ± 10.6 | 81.54 ± 9.18 | 88.14 ± 9.25 | 74.01 ± 11.0 |
| monkey M |  |  |  |  |
| 2 buttons | 87.67 ± 4.66 | ... | 86.42 ± 7.16 | 82.33 ± 3.68 |

**S4 Figure.**
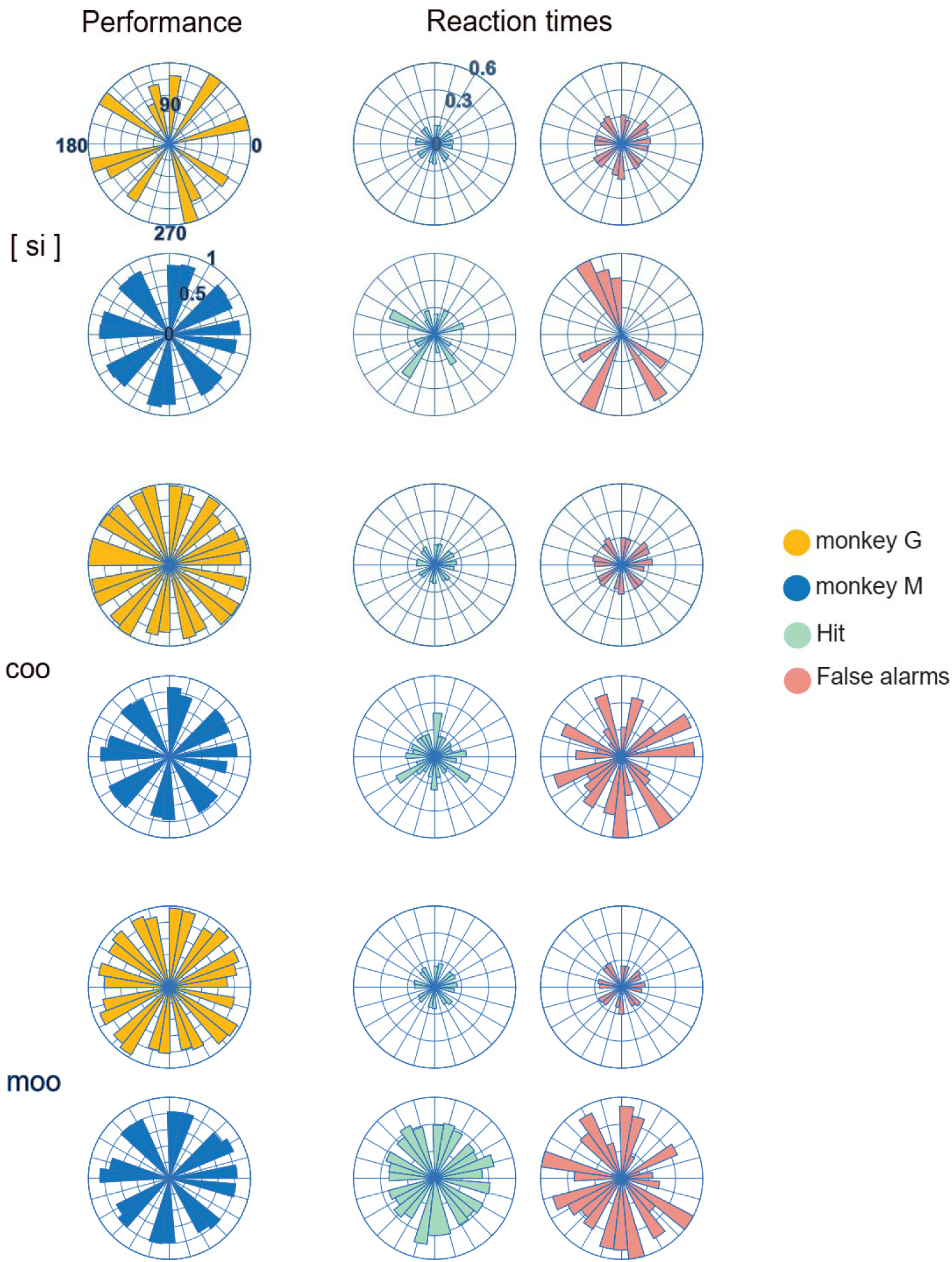
Monkeys’ performance in numerous positions of P on the touchscreen. To assess whether monkeys associated categories with specific regions on the screen, we analyzed the performance of each CC as a function of the angles at which P was presented [one-way ANOVA, False Discovery Rate (FDR) corrected for multiple comparisons using Benjamini-Hochberg procedure (critical p-value = 0.0143)]. In monkey M, we did not find any bias toward a particular angle ([si]: F (11, 4.70215) = 1.95, p = 0.034; [coo]: F (23, 30.6495) = 1.57, p = 0.044; [moo]: F (23, 28.1022) = 0.88, p = 0.632). In monkey G, the performance of the [si] category was not affected by P angles (F (15, 130.086) = 0.36; p = 0.988), but we found a significant effect for the categories [coo] (F(15, 160.67) = 1.97; p = 0.014) and [moo] (F(15, 150.619) = 2.51; p = 0.001). To make sense of these angle effects, we divided the pairwise multiple comparisons (Tukey’s HSD, p<0.05) into quadrants. We found such differences between angles close to each other (<90°) and distributed across all the screen quadrants. In other words, besides significant differences in the performance as a function of the visual Target’s angle, it did not show a systematic quadrant bias. With these finding, all experiments presented here include presentations of pictures in only four quadrants. **S5 Video 2**. Video of monkey M performing in a control condition of the CME task.

**Table 3.** S6 Mean and standard deviation of the performance across all sessions (days). **S6 Table Versions**. Table of the performance of the monkeys during presentations of different versions of the learned sounds.

| Monkey G | coo | moo | ['ro.xo] | [si] |
| --- | --- | --- | --- | --- |
| o | 91.4 ± 16.98 | 84.59 ± 17.17 | 70.34 ± 23.46 | 89.38 ± 19.73 |
| S1 | 73.59 ± 22.31 | 75.99 ± 19.91 | 63.92 ± 21.21 | 89.38 ± 19.51 |
| S2 | 64.23 ± 23.17 | 79.3 ± 19.22 | 62.84 ± 22.19 | 91.18 ± 19.34 |
| S3 | 55.79 ± 26.38 | 72.96 ± 19.84 | 62.62 ± 21.09 | 90.42 ± 19.69 |
| S4 | 70.07 ± 23.4 | 77.83 ± 19.9 | 65.91 ± 19.34 | 91.2 ± 19.77 |
| S5 | 80.98 ± 21.23 | 77.58 ± 17.52 | 72.23 ± 22.33 | 74.86 ± 24.72 |
| S6 | 82.97 ± 20.29 | 70.16 ± 20.97 | 72.52 ± 19.97 | 71.94 ± 24.87 |
| S7 | 89.27 ± 17.21 | 52.47 ± 21.19 | 69.89 ± 21.94 | 83.15 ± 22.23 |
| S8 | 56.84 ± 25.24 | 73.6 ± 21.32 | 63.62 ± 22.86 | 79.3 ± 21.7 |
| S9 | 24.18 ± 19.36 | 62.95 ± 21.22 | 52.13 ± 24.95 | 90.86 ± 19.44 |
| S10 | 78.17 ± 22.87 | 88.37 ± 15.69 | 65.95 ± 21.59 | 88.77 ± 21.64 |
| o-o | 81.29 ± 9.15 | 87.91 ± 7.9 | 82.04 ± 8.88 | 82.51 ± 9.15 |
| S1-S1 | 79.35 ± 10.73 | 87.2 ± 7.29 | 82.23 ± 8.27 | 81.09 ± 9.15 |
| S2-S2 | 77.5 ± 11.46 | 81.04 ± 11.45 | 86.06 ± 7.64 | 80.49 ± 10.39 |
| S3-S3 | 79.45 ± 9.65 | 85.09 ± 10.29 | 82.81 ± 10.07 | 81.45 ± 9.05 |

| Monkey M | coo | ['ro.xo] | [si] |
| --- | --- | --- | --- |
| o | 92.22 ± 5.91 | 85.83 ± 11.5 | 88.08 ± 12.18 |
| S1 | 74.01 ± 19.01 | 77.05 ± 16.61 | 72.78 ± 21.95 |
| S2 | 87.83 ± 11.91 | 57.5 ± 18.12 | 60 ± 25.5 |
| S3 | 78.45 ± 17.01 | 87.21 ± 11.4 | 56.11 ± 21.76 |
| S4 | 83.81 ± 13.02 | 86.25 ± 8.67 | 65.99 ± 25.59 |
| S5 | 70.24 ± 19.87 | 47.5 ± 16.47 | 67.22 ± 19.64 |
| S6 | 88.75 ± 9.91 |  |  |

## References

1. Adachi I, Hampton RR (2011) Rhesus monkeys see who they hear: Spontaneous cross-modal memory for familiar conspecifics. PLoS One 6.

2. Arcaro MJ, Schade PF, Vincent JL, Ponce CR, Livingstone MS (2017) Seeing faces is necessary for face-patch formation HHS Public Access Author manuscript. Nat Neurosci 20:1404–1412.

3. Beauchamp MS, Lee KE, Argall BD, Martin A (2004) Integration of Auditory and Visual Information about Objects in Superior Temporal Sulcus. Neuron 41:809–823.

4. Bodin C, Belin P (2020) Exploring the cerebral substrate of voice perception in primate brains. Philos Trans R Soc B Biol Sci 375.

5. Bodin C, Trapeau R, Nazarian B, Sein J, Degiovanni X, Baurberg J, Rapha E, Renaud L, Giordano BL, Belin P (2021) Functionally homologous representation of vocalizations in the auditory cortex of humans and macaques. Curr Biol 31:4839–4844.e4.

6. Calvert G a, Hansen PC, Iversen SD, Brammer MJ (2001) Detection of audio-visual integration sites in humans by application of electrophysiological criteria to the BOLD effect. Neuroimage 14:427–438.

7. Chandrasekaran C, Lemus L, Ghazanfar AA (2013) Dynamic faces speed up the onset of auditory cortical spiking responses during vocal detection. Proc Natl Acad Sci U S A 110:E4668–77.

8. Cohen YE, Theunissen F, Russ BE, Gill P (2007) Acoustic features of rhesus vocalizations and their representation in the ventrolateral prefrontal cortex. J Neurophysiol 97:1470– 1484.

9. Diehl MM, Plakke BA, Albuquerque R, Romanski LM (2022) Representation of Expression and Identity by Ventral Prefrontal Neurons. Neuroscience 496:243–260.

10. Fuster JM, Bodner M, Kroger JK (2000) Cross-modal and cross-temporal association in neurons of frontal cortex. Nature 405:347–351.

11. Gaffan D, Harrison S (1991) Auditory-visual associations, hemispheric specialization and temporal-frontal interaction in the rhesus monkey. Brain 114:2133–2144.

12. Hickok G, Poeppel D (2007) The cortical organization of speech processing. Nat Rev Neurosci 8:393–402.

13. Huang Y, Brosch M (2016) Neuronal activity in primate prefrontal cortex related to goal-directed behavior during auditory working memory tasks. Brain Res 1640:314–327.

14. Jordan KE, Brannon EM, Logothetis NK, Ghazanfar AA (2005) Monkeys match the number of voices they hear to the number of faces they see. Curr Biol 15:1034–1038.

15. Kawahara H, Masuda-Katsuse I, De Cheveigné A (1999) Restructuring speech representations using a pitch-adaptive time–frequency smoothing and an instantaneous-frequency-based F0 extraction: Possible role of a repetitive structure in sounds. Speech Commun 27:187–207.

16. Khandhadia AP, Murphy AP, Romanski LM, Bizley JK, Leopold DA (2021) Audiovisual integration in macaque face patch neurons. Curr Biol 31:1826–1835.e3.

17. Lai C, Tanaka S, Harris TD, Lee AK (2023) Volitional activation of remote place representations with a hippocampal brain-machine interface. Science 382:566–573.

18. Leaver AM, Rauschecker JP (2010) Cortical representation of natural complex sounds: effects of acoustic features and auditory object category. J Neurosci 30:7604–7612.

19. Lemus L, Hernández A, Luna R, Zainos A, Romo R (2010) Do sensory cortices process more than one sensory modality during perceptual judgments? Neuron 67:335–348.

20. Lemus L, Lafuente V De (2022) Look Who is Talking. Identities and Expressions in the Prefrontal Cortex. Neuroscience 496:241–242.

21. Leopold DA, Bondar I V., Giese MA (2006) Norm-based face encoding by single neurons in the monkey inferotemporal cortex. Nature 442:572–575.

22. Melchor J, Vergara J, Figueroa T, Morán I, Lemus L (2021) Formant-Based Recognition of Words and Other Naturalistic Sounds in Rhesus Monkeys. Front Neurosci 15:1–10.

23. Morán I, Perez-Orive J, Melchor J, Figueroa T, Lemus L (2021) Auditory decisions in the supplementary motor area. Prog Neurobiol 202:1–11.

24. Ng CW, Plakke B, Poremba A (2009) Primate auditory recognition memory performance varies with sound type. Hear Res 256:64–74.

25. Nieder A (2012) Supramodal numerosity selectivity of neurons in primate prefrontal and posterior parietal cortices. Proc Natl Acad Sci 109:11860–11865.

26. Noesselt T, Rieger JW, Schoenfeld MA, Kanowski M, Hinrichs H, Heinze HJ, Driver J (2007) Audiovisual temporal correspondence modulates human multisensory superior temporal sulcus plus primary sensory cortices. J Neurosci 27:11431–11441.

27. Ohayon S, Freiwald WA, Tsao DY (2012) What Makes a Cell Face Selective? The Importance of Contrast. Neuron 74:567–581.

28. Parise C V., Spence C, Deroy O (2016) Understanding the correspondences: Introduction to the special issue on crossmodal correspondences. Multisens Res 29:1–6.

29. Ratcliffe VF, Taylor AM, Reby D (2016) Cross-Modal Correspondences in Non-human Mammal Communication. Multisens Res 74:49–91.

30. Ravignani A, Sonnweber R (2017) Chimpanzees process structural isomorphisms across sensory modalities. Cognition 161:74–79.

31. Rescorla RA, Wagner AR (1972) A theory of Pavlovian conditioning: Variations in the effectiveness of reinforcement and non-reinforcement. In: In Classical conditioning II (Black A, Prokasy W, eds), pp 64–99. New York: Appleton-Century Crofts.

32. Romanski LM (2007) Representation and integration of auditory and visual stimuli in the primate ventral lateral prefrontal cortex. Cereb Cortex 17:61–69.

33. Scott BH, Mishkin M, Yin P (2012) Monkeys have a limited form of short-term memory in audition. Proc Natl Acad Sci U S A 109:12237–12241.

34. Seyfarth RM, Cheney DL, Marler P (1980) Vervet monkey alarm calls: Semantic communication in a free-ranging primate. Anim Behav 28:1070–1094.

35. Stephen EP, Li Y, Metzger S, Oganian Y, Chang EF (2023) Latent neural dynamics encode temporal context in speech. Hear Res 437:1–13.

36. Town SM, Wood KC, Bizley JK (2018) Sound identity is represented robustly in auditory cortex during perceptual constancy. Nat Commun 9:1–15.

37. Treviño M (2016) Associative learning through acquired salience. Front Behav Neurosci 9:168673.

38. Tsao DY, Freiwald W a, Knutsen T a, Mandeville JB, Tootell RBH (2003) Faces and objects in macaque cerebral cortex. Nat Neurosci 6:989–995.

39. Tsunada J, Lee JH, Cohen YE (2011) Representation of speech categories in the primate auditory cortex. J Neurophysiol 105:2634–2646.

40. Tyree TJ, Metke M, Miller CT (2023) Cross-modal representation of identity in the primate hippocampus. Science 382:417–423.

41. Walker L, Walker P, Francis B (2012) A common scheme for cross-sensory correspondences across stimulus domains. Perception 41:1186–1192.

42. Weiskrantz L, Cowey A (1975) Cross modal matching in the rhesus monkey using a single pair of stimuli. Neuropsychologia 13:257–261.

43. Yi HG, Leonard MK, Chang EF (2019) The Encoding of Speech Sounds in the Superior Temporal Gyrus. Neuron 102:1096–1110.

44. Zhou YD, Fuster JM (2000) Visuo-tactile cross-modal associations in cortical somatosensory cells. Proc Natl Acad Sci U S A 97:9777–9782.

